# SMART Sensors for Cell Growth Monitoring in Closed System G-Rex for Scalable, Cost-Conscious Cell and Gene Therapy Manufacturing

**DOI:** 10.1101/2025.09.17.676768

**Authors:** Subhanwit Roy, Samuel Rothstein, Aidan Shervheim, Yee Jher Chan, Cameron Greenwalt, Nathan Neihart, Jose Juarez, Patricia Turner, Dan Welch, John Wilson, Dan Fick, Nigel F. Reuel

**Affiliations:** Skroot Laboratory Inc. – Ames, Iowa; CellReady LLC – Houston, Texas; Wilson Wolf Manufacturing LLC – St. Paul, Minnesota

**Author notes:** Corresponding Author –.

**Keywords:** Process analytical technologies (PAT), cell expansion, contact free, static bioreactor, cell quantification

## Abstract

Cell and gene-modified cell therapy (CGT), commonly referred to as the fourth pillar of cancer treatment, includes creating advanced cancer killing cells *ex vivo* and then subsequently putting them into the patient’s body to selectively attack cancerous cells. Typically, immune cells from a patient’s body are removed, genetically modified to find and kill cancer, grown to a large army of cancer killing cells, and put back into the patient. This complex process is being simplified through innovation, some of which we describe here. One key innovation is an immune cell manufacturing device configuration called G-Rex (for “gas permeable rapid expansion”), which is now widely used to create armies of cancer killing cells for clinical trials and for commercially approved drug products on a worldwide basis. G-Rex is uniquely simple and popular because it is the only technology that eliminates the need for pumps and flow circuits to deliver oxygen and nutrients. Instead, the material and geometry of G-Rex allow cancer killing cells to get access to oxygen and nutrients on demand. This greatly minimizes complexity and sets the stage for high throughput manufacturing.

To further simplify G-Rex relative to all CGT drug product manufacturing alternatives, it would be ideal to eliminate any need of physical intervention to assess cell quantity. With that goal, we conceived a novel approach to determining cell quantity in a non-invasive manner. More specifically, we invented and built a sensor into G-Rex that continuously reports the quantity of cells in G-Rex without any intervention or sampling to an external reader. Herein, we call the sensor a “single-use metabolite absorbing resonant transducer in G-Rex (SMART G-Rex)” and the reader a “SMART Reader.” The SMART G-Rex is wirelessly interrogated by the SMART Reader to determine the extent of cell expansion in the G-Rex at any given time. More specifically, as cell quantity increases, the SMART G-Rex absorbs cell secreted metabolites (organic compounds such as eicosanoids) which cause a change in resonant frequency (measured as % of frequency change, or Skroot Growth Index (SGI)). By placing SMART G-Rex in proximity of the SMART Reader, whether in or out of an incubator, changes in resonant frequency are detected and translated into cell quantity with high reliability. For example, using 45 sensors across multiple donor cell samples, we established 99% confidence in SMART’s accuracy at signaling a >60X expansion (3B cells at harvest). The SMART G-Rex provides a non-invasive way of instantly quantifying the number of cells in a CGT drug production process and informing manufacturers when a CGT drug product has reached the desired quantity of cells necessary to proceed with formulation, fill, and finish. By integrating SMART technology, G-Rex is the only technology for CGT manufacturing that can avoid intervention when determining cells quantity or allow real-time knowledge of cell quantity.

**Graphical Abstract:** 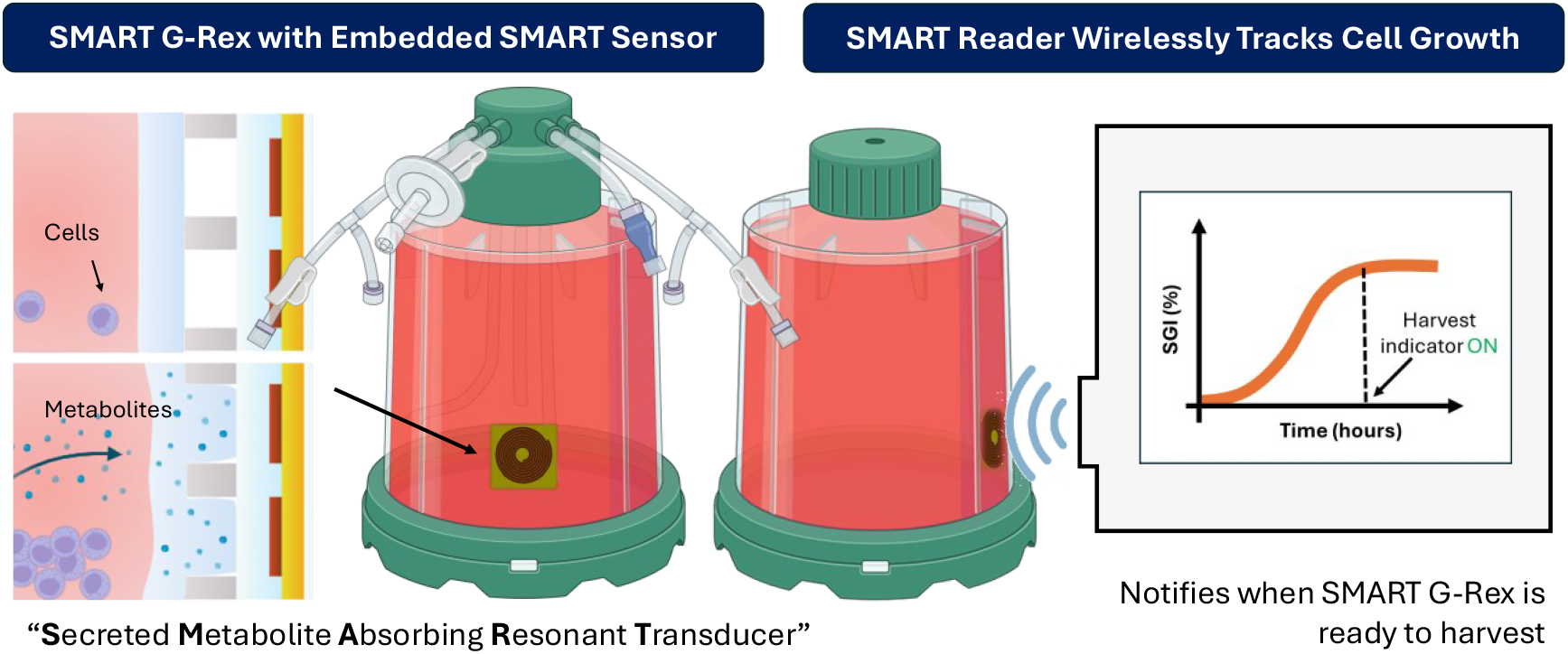

## Introduction

Personalized cell therapies are opening a new epoch in drug development [1]. Chimeric antigen receptor T (CAR-T) cell therapies, in which a patient’s T-cells are genetically engineered to express a CAR that is designed to recognize specific tumor targets, have become mainstream cancer treatments [2]. Seven CAR-T drug products are commercially available for leukemia, lymphoma, and myeloma and over 1000 clinical trials being pursued globally for cancer and more recently for autoimmune diseases [3]. The treatable patient population continues to rise. One estimate identified 450,000 eligible patients in 2019, with projected growth to 2 million in 2029 [4]. But the capacity to manufacture drug products for this number of people is in question. Manufacturing limitations are made clear by the fact that as of 2025, barriers to manufacturing scale out are being reported by commercial drug manufacturers [5–8].

A root cause of manufacturing limitations includes the fact that CAR-T drug products typically originate in academic centers where state-of-the-art manufacturing skills are of little importance compared to knowledge of the mechanisms of action in a drug product that drive safety and efficacy. This results in wide differences in every element of the manufacturing process from drug product to drug product and academic institute to academic institute. As the private sector spins these drug products out of the academic centers for commercialization, the poorly conceived manufacturing processes follow. From there, companies focus on quickly gaining more clinical data, which takes precedent over stopping to improve the manufacturing process. With each company chasing clinical data and operating in a silo, there has been little focus on standardizing elements of the drug product manufacturing process. Yet, there is ample opportunity to standardize manufacturing because the manufacturing steps of every CGT drug product have much in common [9]. At a minimum, three steps are certain to be present. Cells need oxygen, cells need nutrients, and the quantity of cells needs to be known before manufacturing is terminated to ensure the patient dose requirements are met.

The simplest technology to provide cells with oxygen and nutrients is called G-Rex which stands for Gas Permeable Rapid Expansion of cells [10]. When cells are in G-Rex, they gravitate through nutrients containing medium to the bottom of G-Rex where they reside on a highly gas permeable membrane and as cells consume oxygen, they create an oxygen concentration gradient across the gas permeable membrane. In response, oxygen from external air diffuses into G-Rex at the rate cells need it. At the same time, G-Rex holds a large volume of nutrients that feed cells by naturally occurring convective movement as cells need it.

Another key attribute of G-Rex is that it occupies very little space and can use standard cell culture incubators for temperature and pH control. Absent oxygen and nutrient pumping mechanisms that are inherent to all other forms of CGT drug product manufacturing processes, G-Rex is unencumbered by the space requirements of pumping equipment and about 40 G-Rex devices can occupy a standard cell culture incubator [11]. On the contrary, competing pump-based technology requires its own custom incubator. The net result is that on a space occupancy metric, G-Rex in comparison to pumping systems can produce drug products at a ratio of about 40 patients to just 1. **Figure 1** provides a space comparison of G-Rex with a typical system that force feeds cells with oxygen and nutrients.

**Figure 1.**
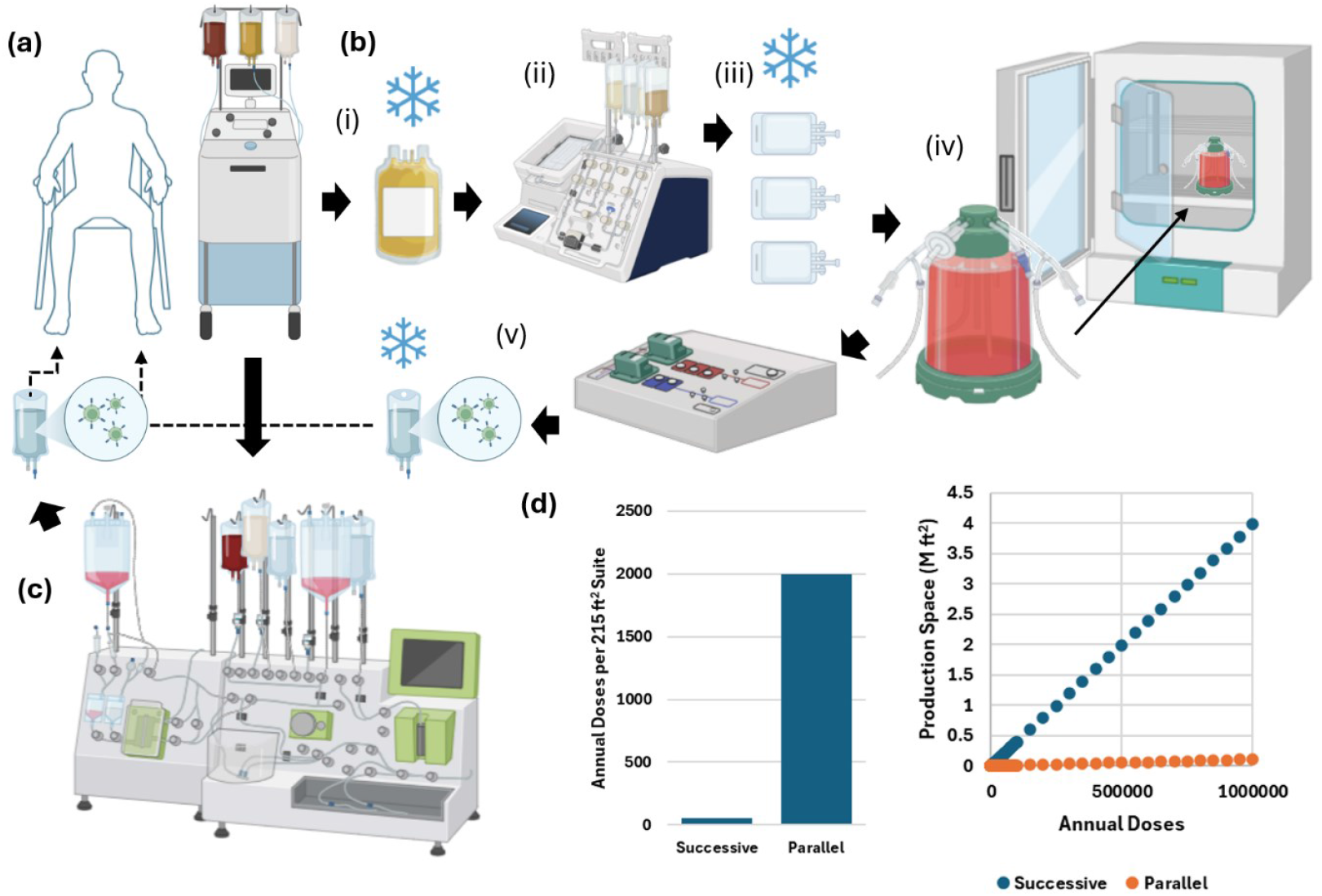
Comparison of the parallel CAR-T process (assembly line with G-Rex) and successive single dose per unit processes. In both workflows, patient cells are collected and separated via leukapheresis (a). (b) CAR-T production with G-Rex has discrete, parallelizable unit operations: i) frozen leukocytes are thawed; ii) T cells are enriched and then aliquoted (iii) to individual dose size; (iv) these are dosed into G-Rex vessels, activated, transduced, and expanded; (v) cells are then harvested and stored as a patient dose. (c) Illustration of a successive, single dose system, such as the CliniMACS Prodigy, that has limited, serial production. (d) Comparison of annual throughput for both systems and scaling of production floor space required.

This uniquely simple method of oxygenating and feeding cells in G-Rex maximizes the utilization of space, personnel, and equipment which reduces cost of goods sold (COGS). But more importantly, this simple approach to manufacturing allows a level of scalability that is unattainable when cells are force fed with oxygen and nutrients. The net result is lowered manufacturing cost and greater patient access.

Despite all the advantages of G-Rex for manufacturing CGT drug products, like all other CGT manufacturing technology, it still carries a need for intervention to determine cell quantity. Cells expand best when left undisturbed on the gas permeable membrane, and thus traditional cell counting via resuspension and sample collection is not recommended during the expansion phase in the G-Rex. Instead, the current best practice is to sample the media, leaving the cells undisturbed, and measure the reduction in glucose and/or increase in lactate levels to monitor cell growth progression [12]. This sampling methodology requires careful handling of each G-Rex and unique or custom-made consumables to facilitate closed system sampling, or a biosafety cabinet to remove the samples in an open system and then run them through an analyzer, most frequently test strips. After expansion is completed (lactate levels plateau) the cells are harvested, and a cell count is used to confirm cell quantities sufficient for patient dosing have been achieved. Each of these hands-on steps contribute to the direct labor costs of the final product. We have discovered a way to eliminate the need for these steps through integration of the SMART sensor.

We now describe the application of single use metabolite absorbing resonant transducer (SMART) sensor, built into G-Rex, that informs the user of cell quantity without any intervention. The inherent benefit of the SMART sensor as an alternative to human intervention is to reduce direct labor cost, eliminate the risk of contamination, and allow reliable monitoring of cell growth without any need to open the G-Rex or remove from an incubator for such assessment.

## Methods

1. Sensor fabrication and testing – Each sensor was 1 inch (2.54 cm) square and consisted of five layers (from fluid-facing side to vessel-facing side): a transduction membrane responsive to secondary metabolites, a polyethylene terephthalate (PET) sheet with engineered voids, an adhesive tie layer, a resonator (copper coil on polyimide), and an adhesive backing for vessel attachment (**Fig. 2a**). The resonator at the core of each sensor was first patterned by chemically etching a copper-clad polyimide laminate. To enable batch fabrication, multiple resonators were etched on a single sheet. Sensors were then hand-assembled as a multilayer stack: the vessel-facing adhesive was applied to the polyimide side of the resonator sheet, followed by application of the adhesive tie layer to the copper side, and alignment of the PET sheet onto the tie layer. This partial stack was laminated using a cold press to remove air pockets and ensure bonding. Finally, the transduction membrane was added, and the full assembly was laminated again to produce a smooth, bubble-free sheet. Individual 2.54 cm square sensors were later cut from this stacked sheet. To place the sensor inside the G-Rex vessel, the backing liner of the vessel-facing adhesive was removed, and the sensor was loaded into a custom-designed positioning guide. This guide ensured consistent placement within the vessel – both in terms of distance from the gas-permeable membrane and radial sector location. Once inserted, a squeegee was used to press the sensor firmly onto the interior vessel wall. After sensor placement, the liner protecting the transduction membrane was removed, and the G-Rex was assembled following Wilson Wolf’s standard assembly procedure. The transduction method has been described in prior work [13]. In brief, as cells grow, they secrete secondary metabolites (organic communication compounds) that absorb selectively into the SMART sensor membrane and act as a plasticizer, softening the membrane material to fill engineered air gaps in the sensor structure (**Fig 2b**). In our prior screening work, we observed simple terpenoids, such as prenol, caused a strong sensor response. In the case of T cells, there is evidence of secreted lipid mediators, specifically eicosanoids and their derivatives, such as prostanoids and leukotrienes, as signaling molecules [14]. We have verified that arachidonic acid (AA), an omega-6 PUFA that is a precursor to eicosanoids, causes a strong sensor response (Supplement 1). This is representative of a broad class of organic compounds that these cells secrete during active growth which cause the sensor to respond. A key advancement over prior work in the SMART G-Rex was the use of laser cut PET for the structured air voids allowing for larger-scale sensor production and greater consistency over the previous wire wound sensors. When the membrane softens and the air voids are filled, the growth media is physically closer to the resonant coil causing a large increase in local permittivity, increasing the capacitance of the coil, resulting in a measurable decrease in the sensor’s resonant frequency. Additionally, increasing the width of the PET air gaps (kerf) emerged to be a useful design parameter for tuning sensor sensitivity (Supplement 2). Sensor consistency was measured by placing replicates (n=4) sensors on the same vessel and determining the coefficient of variation in the sensor response. Effects of scaling production were also determined by outsourcing the production to a flexible circuit manufacturer (Flexible Circuit Technologies) and comparing to in-house assembled sensors. The third-party sensors were further subjected to cytotoxicity testing and accelerated aging to determine the robustness of this method as a commercial product. Cytotoxicity testing of the sensor placed in a polycarbonate cap was executed at NAMSA to evaluate for potential cytotoxic effects using an *in vitro* mammalian cell culture test according to ISO 10993-5 and BS EN ISO 10993-5, Biological evaluation of medical devices - Part 5: Tests for in vitro cytotoxicity, whereas accelerated aging was performed at Skroot by placing vessels with attached sensors in a Fisherbrand Oven at 59 °C for 4, 8, and 12 weeks to achieve equivalent aging of 1, 2, and 3 years respectively [15], following which cell culture tests were performed at CellReady, LLC to evaluate sensor response of the differently-aged vessels (Supplement 3).
2. Reader design – The resonant frequency of the sensor was wirelessly interrogated through the vessel using a custom-built reader that performed frequency sweeps across a defined frequency range. Two reader configurations were implemented using the same core platform: 1) a continuous reader mode placed inside an incubator (**Fig. 2c**), interrogating the sensor every 5 minutes, and 2) a bench-top reader mode (**Fig. 2d**) in which the G-Rex was removed from the incubator, placed on the reader, and scanned for a single reading in under 30 seconds. The design and operation of the reader platform have been detailed in previous work [16]. Briefly, the reader applies a sinusoidal excitation (the frequency of the sinusoid can be digitally controlled) to an inductive loop that electromagnetically couples with the sensor, and the voltage measured across this loop yields a signal that exhibits a peak at the sensor’s resonant frequency, which reflects changes in the sensor’s local environment. This design improved on prior work to incorporate a larger frequency sweep range and advanced filtering of the output voltage signal. Custom software was used to extract the resonant frequency from the voltage signal (Supplement 4). To reduce systematic error, a software-driven calibration step was performed using a reader-only measurement (with no sensor present) to account for the reader’s intrinsic frequency response. The final sensor output was reported as the Skroot Growth Index (SGI; see **Fig. 2e**), defined as the percentage change in resonant frequency from a pre-determined equilibration time (e.g., 24 hours after the start of the run):

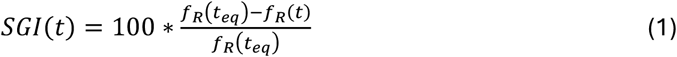

where *f_R_* denotes the sensor resonant frequency and *t_eq_* represents the equilibration time. The use of a finite *t_eq_* allowed for the elimination of any initial drift in sensor response caused by mechanical effects, ensuring that subsequent changes in SGI are primarily driven by biological phenomena.
3. Cell experiments – To validate the sensor response for use in a G-Rex based CAR-T manufacturing process, extensive cell culture testing was done at CellReady, LLC using cryopreserved Peripheral Blood Mononuclear Cells (PBMCs) as the starting material. To evaluate the sensors’ ability to track cell growth for different growth profiles indicative of biological differences, three standard growth conditions were established by varying the seeding density: fast (200 million activated cells), medium (50 million activated cells), and slow (20 million activated cells). Following CellReady’s standard protocol for activating and expanding T cells from PBMCs, a bulk quantity of PBMCs were thawed, rested, and activated for 48-72 hours in a single vessel, prior to seeding the activated cells for the sensor experiments in multiple G-Rex 100M’s. Subsequently, activated cells were seeded into G-Rex 100M devices with SMART sensors using Human T Cell media, according to the assigned growth condition. Reader measurements were initiated upon seeding and cells were allowed to expand for a pre-determined period ranging from 4 to 8 days. Standard reference measurements of glucose and lactate were also performed on a daily basis starting 4 days after seeding by carefully opening the G-Rex in a sterile environment, removing a 100 µL sample, and measuring with handheld meters – ForaCare Inc. FORA G20 Blood Glucose Monitor was used for glucose measurements whereas Hearts Bio Inc. Heartscare C1 lactate meter and Nova Biomedical Lactate Plus meters were used for lactate measurements. Upon conclusion of each cell culture run, the cells were harvested, and the total cell count, and final cell viability were assessed using a Cellometer K2 Fluorescent Cell Counter by Nexcelom (now a part of Revvity).
4. Data Collection and Harvest algorithm – Based on the data provided by the reader, SGI from the equilibration time (corresponding to SGI = 0) was determined by the software. In the case of the continuous read, this was done by referencing the pre-set equilibration time. In benchtop mode, the vessel was tracked by a unique identifier (barcode) allowing comparison between scan points. The software allowed for tracking the change in SGI at each data point, determining when the sensor response slowed and eventually plateaued. This plateau point was then used to correlate to cell growth data to determine trigger points for harvesting messaging.

**Figure 2.**
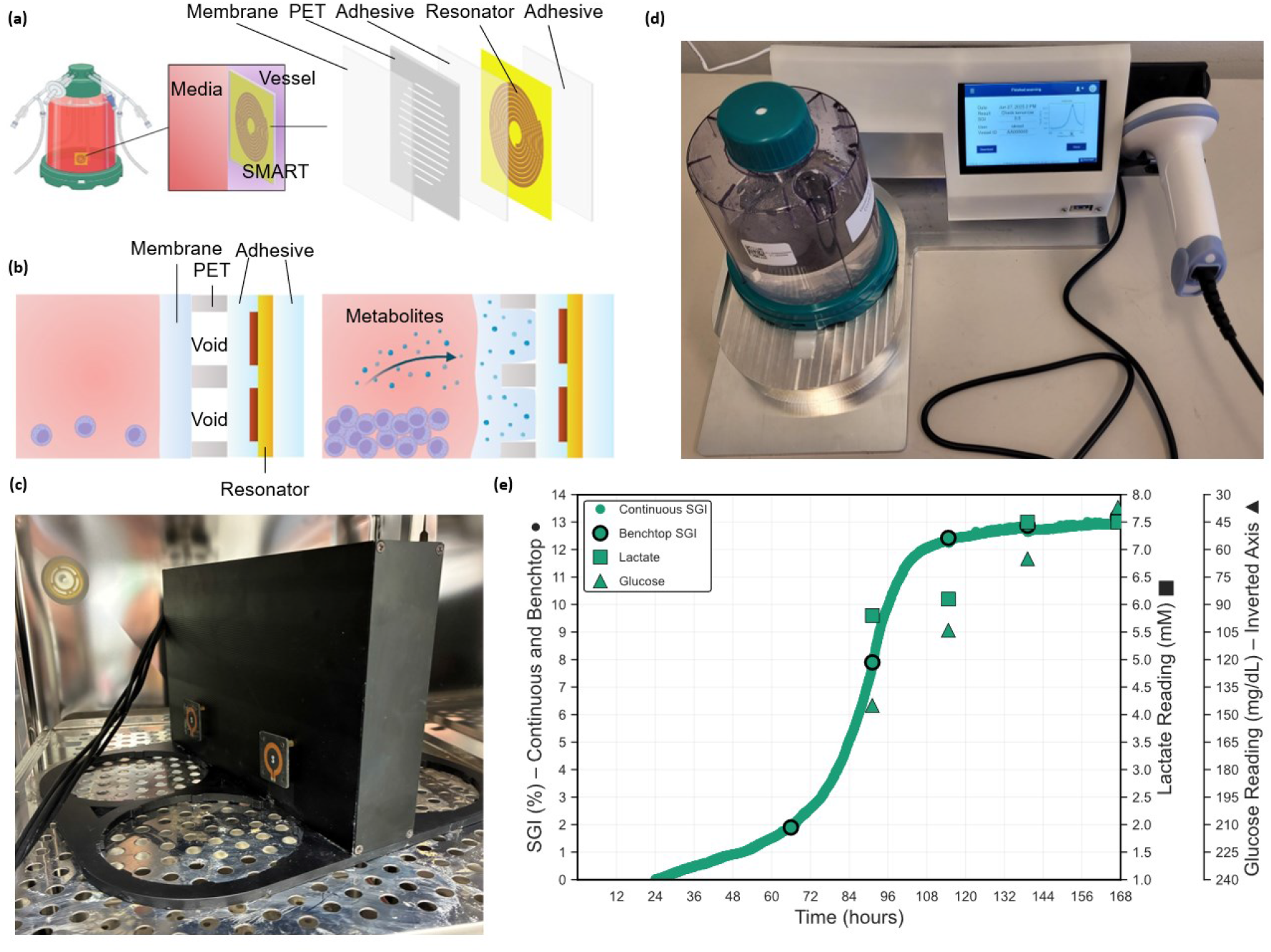
SMART sensor structure and response. (a) The SMART sensor sticker is adhered to the inside of the G-Rex vessel to be in contact with the growth medium and is comprised of five layers (from fluid side: responsive membrane, PET sheet with engineered voids, adhesive tie layer, resonator, and adhesive layer for bonding to vessel interior. (b) Sensor transduction mechanism: as the cells secrete secondary metabolites (terpenoids) the membrane softens, fills the voids and the local permittivity to the resonant coil changes, causing a change in resonant frequency. (c) Custom reader to interrogate the sensors continuously within an incubator. (d) Custom reader to interrogate the sensors intermittently in a bench-top reader mode. In both cases, the G-Rex is kept closed and the cell layer is undisturbed. (e) Response of sensor reported as the change in resonant frequency from the start point (Skroot Growth Index, SGI) in both continuous and benchtop mode compared to traditional

## Results and Discussion

### Sensor Performance

Initial testing of the sensor in continuous reader mode used a commercial portable vector network analyzer (VNA) as the reader hardware. It showed marked changes in cell production versus control vessels. Upon inspection, increased condensation was observed in the continuous read vessels and confirmed by temperature monitoring that indeed the continuous reader had raised the G-Rex temperature by ~1 °C, adversely affecting cell growth rate (Supplement 5). This was overcome by custom-designing the reader with special attention to power consumption of selected electronics parts and programming it to enter sleep mode between measurements, thereby reducing the overall heat generation and maintaining a lower board temperature. Using this cooler hardware, cell growth between vessels with a sensor and controls without a sensor were statistically the same (Supplement 5).

Sensor-to-sensor consistency was assessed by comparing the response of four replicate sensors placed on the same vessel. Because SGI is a derived metric, the underlying measured resonant frequencies were used to quantify consistency via the coefficient of variation (CV) across the full duration of the experiment. For each frequency scan, CV was calculated as the standard deviation divided by the mean of the four sensors [17], expressed as a percentage. **Figs. 3a–c**. show the SGI, measured resonant frequency, and corresponding CV of the four replicate sensors monitoring a CAR-T cell culture from an equilibration time of 24 hours to the end of experiment. The maximum CV remained below 1.25%, indicating high consistency across sensor replicates.

**Figure 3.**
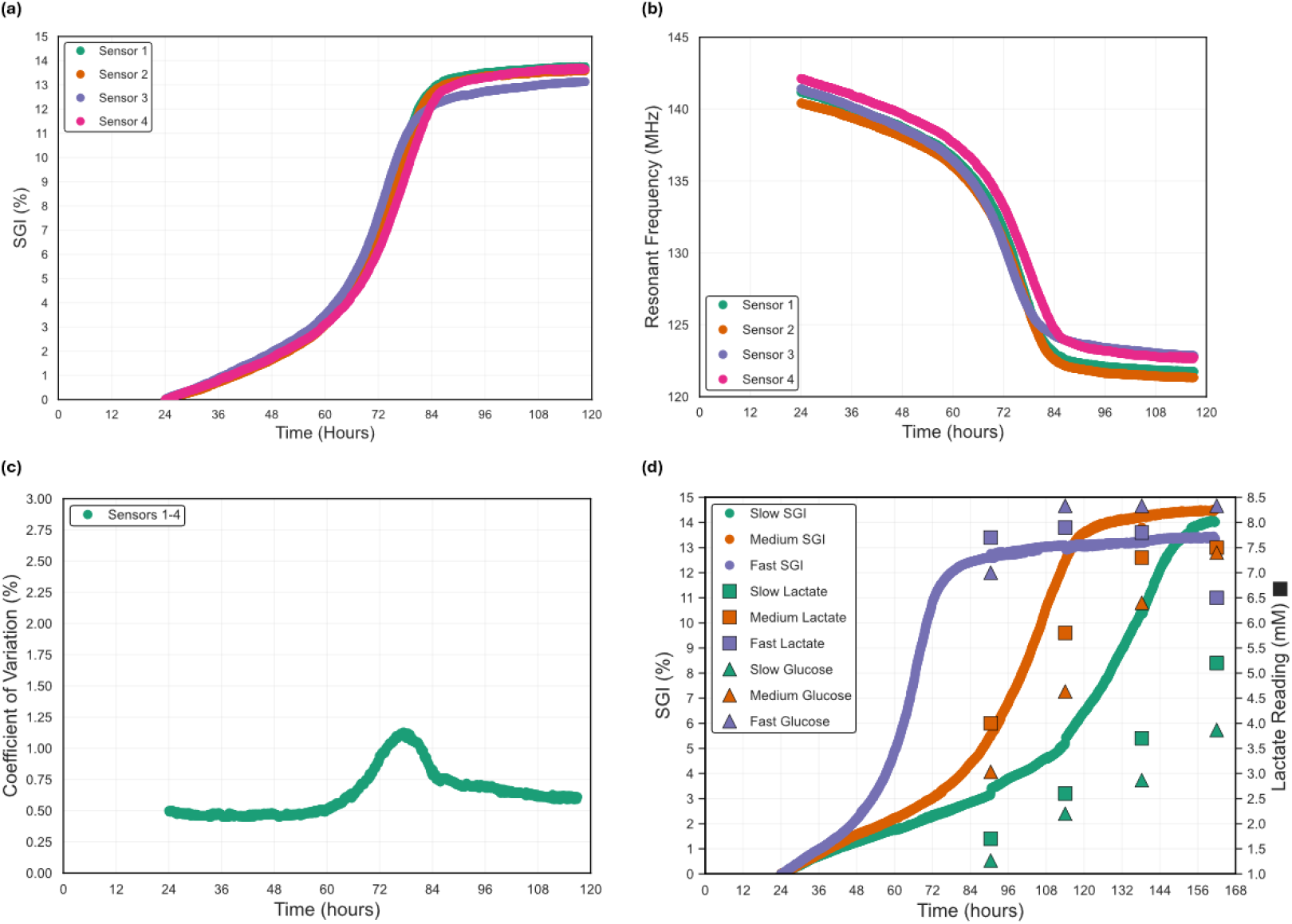
(a) SMART sensor response (Skroot Growth Index – SGI) over time for one G-Rex 100M device with 4 replicate sensors, (b) Corresponding measured resonant frequency data, (c) Consistency among the 4 sensors expressed as coefficient of variation (CV) in measured resonant frequency, (d) SMART sensor response (Skroot Growth Index – SGI) over time for G-Rex 100M devices seeded under slow, medium, and fast growth conditions, alongside manually sampled lactate and glucose measurements. The sensor reliably distinguishes between different cell growth rates, as evidenced by distinct SGI

Having shown sensor consistency, we then evaluated the sensor capability to track different cell growth rates. This was done by seeding 3 independent G-Rex 100M vessels with activated cells from the same donor and the same activation source vessel at the same time with slow, medium, and fast growth conditions, respectively, and recording the sensor response for 7 days (168 hours). The sensor measurements were coupled with daily metabolite profiles in terms of lactate and glucose samples from each vessel, starting Day 4. The SGI data for the three conditions, using an equilibration time of 24 hours, is shown in **Fig. 3d**, clearly highlighting the effectiveness of the sensors in differentially tracking the three growth conditions. It is evident from the correlation between the SGI curves and the lactate and glucose trends that the SGI tracks actual cell growth patterns inside the vessels. Additionally, it was found that the time taken to collect manual sampling parameters for 11 vessels by a GMP specialist was 29 minutes 34 seconds, whereas benchtop reader scans for the same vessels took 14 minutes 45 seconds – resulting in a time-saving of 50% with more improvements possible through software optimization.

Further testing was done to validate sensor response across various cell and vessel types. The Skroot sensor platform proved capable of successfully tracking both bacteria and mammalian cell growth across static and dynamic systems (Supplement 6). Moreover, the sensors demonstrated the ability to readily detect aberrant cell growth patterns. For instance, the presence of biological contaminants in a mammalian cell culture resulted in a markedly accelerated increase in SGI, consistent with the rapid expansion of the contaminants relative to the intended cells (Supplement 6).

### Terminal Cell Count – Verifying Sufficient Expansion Prior to Cryostorage

In standard CAR-T manufacturing, after expansion the cells are counted to verify sufficient dose quantity and viability prior to cryopreservation. The cells are then concentrated and re-suspended in bulk at the concentration specified on the label in the final drug product formulation media including a cryo-preservative and usually some type of Multiple Electrolytes for Injection, Type 1 Solutions, the bulk formulated drug product is then aliquoted into the primary drug product container, which is usually a ethylene-vinyl acetate (EVA) or polyolefin bags and then stored in vapor phase of liquid nitrogen. It is important to note that manufacturers often over-fill primary containers to compensate for the expected loss of cells due to the cryopreservation and thawing process.

Although post-expansion cell counts currently serve as the industry standard for formulating cell therapies at a target concentration, all counting methods inherently carry some degree of variance in total cell counts and recoveries – variability that is widely accepted across the field. Thus, a surrogate technique with a validated variance equal to or smaller than this accepted range could serve as the basis for drug-product formulation. It may be beneficial in some applications to allow a defined dose range on the label, provided a confirmatory count on a thawed sample of the finished product is recorded on the Certificate of Analysis (CoA) with the other quality-control (QC) data.

Taking this concept further, if a manufacturing process generates sufficiently repeatable yields, then formulation processes could be completely standardized on a per-process basis, eliminating the need for operator- or instrument-specific volumetric adjustments. For instance, if the manufacturing process can be validated to generate between 3.0 × 10^9^ to 4.0 × 10^9^total cells in 95% of manufacturing runs, independent of donor starting material, and the SMART sensor can verify that the cell count would fall within this range, then conceivably all the necessary wash buffers and formulation medias could be batch prepared for off the shelf formulation and fill. This approach would greatly simplify formulation, increase throughput, and shorten the interval from harvest to freezing, thereby enhancing process repeatability and improving post-thaw recovery and viability.

The correlation of final SGI saturation levels to final cell counts was explored to explore the feasibility of this concept and determine if this measure could be used reliably to remove the necessity of the post expansion cell count prior to final formulation. We pooled data from 45 sensors that were all seeded at the same cell density (50 million activated cells) but from different donors and were run past SGI saturation. When plotting the final SGI to total viable cells (**Fig. 4a**), there was a weak positive correlation (0.49 Pearson correlation coefficient). This analysis was confounded by the inherent variability of optical based cell counters that are acknowledged to have ±10% accuracy. We also assessed the distribution of final SGI (histogram plotted in **Fig. 4b-c**) and observed a Weibull distribution shape. Most sensors saturated at > 9% SGI for this cell manufacturing process; however, there were a few (n = 3) that had a lower saturation point. By fitting the distributions of the sensor saturation points and the final viable cell counts (histogram and uniform distribution plotted in **Fig. 4d**), we could confirm with 99% confidence that a final SGI > 7.08% corresponds to greater than 3.02 billion cells or a > 60X expansion. Additionally, in vessels where cell growth was intentionally stunted (see DMSO vessels in Supplement 7), SGI saturated at much lower values.

**Figure 4.**
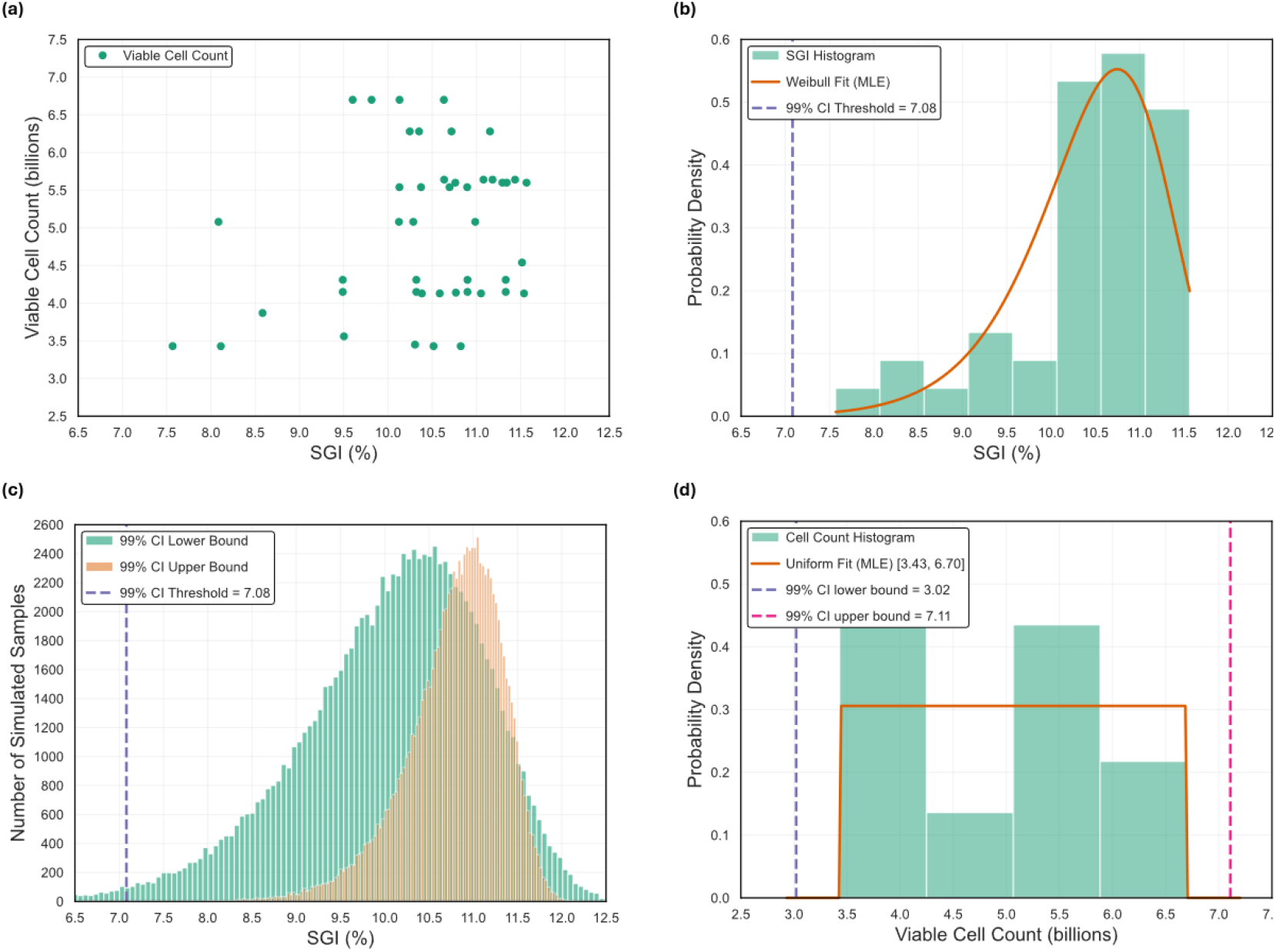
(a) Correlation of total viable cells to final SGI measurement for 45 samples seeded with medium growth condition (50 million cells) exhibiting a weak positive correlation, (b) Histogram of final SGI values for samples overlaid with maximum-likelihood Weibull fit - the dashed purple line marks the 1 % lower‐tail quantile (99 % confidence threshold of 7.08%). (c) Overlaid histograms of 100,000 simulated SGI draws using the lower‐ and upper‐bounds of the 99% confidence interval (CI) on the Weibull parameters. The dashed purple line indicates the 99% “noise‐floor” threshold from panel, (d) Histogram of viable cell counts with maximum-likelihood uniform fit spanning the observed min/max [3.43, 6.70] billion. Dashed purple and pink lines show the analytically derived 99 % CI lower (3.02 billion) and upper (7.11 billion) bounds.

This resulting confidence may be sufficient to eliminate the post expansion cell verification count and simply formulate cells with off the shelf batch prepared formulation reagents and aliquot the harvested cells into five cryopreservation bags (> 600M cells per bag). One of the formulated and finished bags could then be used as the QC verification for final product QC testing, including cell count and viability measurements. If these parameters fall within the labeled range, and all other QC criteria are met, the remaining drug product bags may be released for patient administration. A major advantage of this approach is that it enables QC procedures to be staged and scheduled like a production line, ensuring that all product testing is performed on a representative finished drug product while enabling parallel QC operations on multiple drug product lots. The main caveat is that this methodology requires that the manufacturing process always produces an excess of product. However, given that the dose requirements for the currently approved CAR-T products range from 2 × 10^8^ – 6 × 10^8^ viable CAR+ cells, generating sufficient surplus should be feasible [18]. A standardized G-Rex 100M-CS process will produce greater than 2.0 × 10^9^ CAR+ T cells.

### Harvest Indicator

After confirming the SMART sensor’s ability to track cell growth and investigating how final SGI related with terminal cell count, we evaluated its potential to generate actionable insights by identifying harvest readiness of a cell culture. The objective of determining harvest readiness is to collect cells at peak viable count, which typically occurs during a window shortly after the exponential growth phase. Harvesting too early could result in suboptimal yield, while harvesting too late can lead to a decline in viability as cells begin to die and the overall culture health deteriorates. Due to inherent biological variability of reviving frozen cells as well as varying quality of patient cells, the timing of this harvest window is difficult to predict *a priori*. Therefore, a real-time, sensor-based indicator of harvest readiness would be a valuable tool for improving process consistency and maximizing product quality.

To develop such an indicator, we designed an experiment in which three G-Rex devices were seeded identically with the medium growth condition (50 million cells) on the same day and harvested on Days 5, 6, and 7 post-seeding. Sensor responses from all three vessels were monitored continuously, and the viable cell count from each vessel was assessed upon harvest (**Fig. 5a**). The cell count results show that the Day 5 and Day 6 harvests were premature, yielding less than two-thirds the viable cell count achieved with the Day 7 harvest.

**Figure 5.**
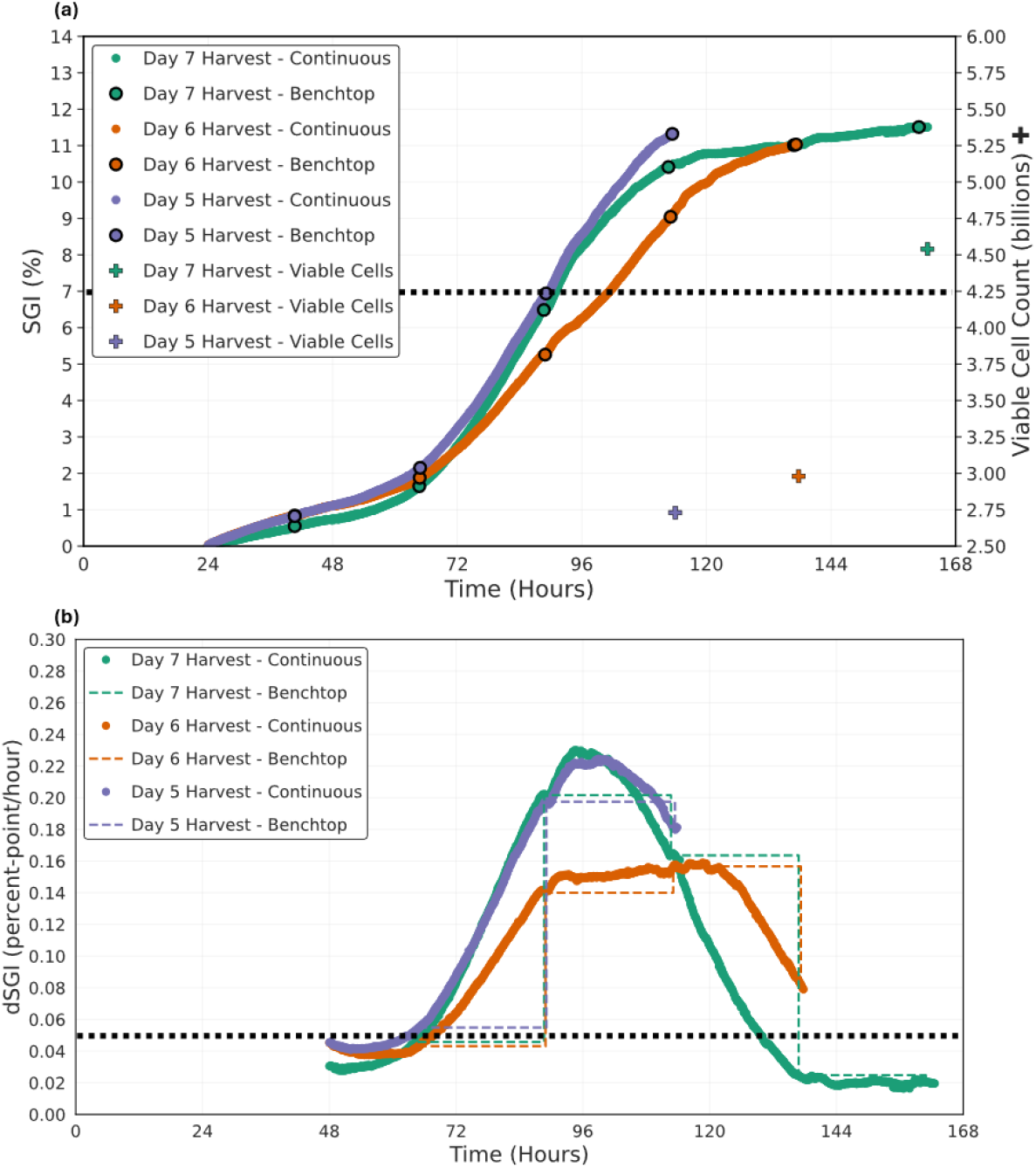
(a) SMART sensor response (Skroot Growth Index – SGI) over time for three G-Rex 100M devices seeded identically with medium growth condition and harvested on three separate days (Day 5, 6, and 7 from seeding). The right Y axis denotes the viable cell count at the time of harvest for each vessel. Discrete data points (black-circled markers) represent benchtop proxy measurements sampled from the continuous SGI curve. The dotted dashed black line denotes the threshold which final SGI needs to cross to have a viable cell count greater than 3 billion. (b) The rate of change of SGI (dSGI) for the continuous measurements and the benchtop proxy. A binary harvest decision rule on dSGI activated upon SGI crossing 7 and using a threshold of 0.05 percent-point/hour for dSGI (dashed black horizontal line) would have indicated that the Day 5 Harvest and Day 6 Harvest vessels were not ready at the time of harvest, while the Day 7 Harvest vessel would have been identified as harvest-ready on Day 6. Notably, harvesting after dSGI dropped below the threshold resulted in a significantly higher viable cell count.

To compare the utility of continuous and benchtop reader modes for harvest decision-making, discrete points from the continuous SGI curves were sampled at each harvest time to serve as proxies for benchtop measurements (**Fig. 5a)**. This approach is valid as both reader modes use the same underlying operating mechanism, and benchtop reader measurements indeed correspond to discrete samples on the continuous SGI curve (see **Fig. 2e**). Additional benchtop proxy measurements were generated at 24-hour intervals prior to the first harvest.

The numerical derivatives of both the continuous and benchtop SGI (denoted as dSGI) were computed for all three vessels (**Fig. 5b**). While the continuous dSGI plots exhibited a bell-curve like response, the benchtop dSGI value exhibited expected stepwise behavior. After the SGI value crossed 7% (approximate SGI value beyond which saturatioguarantees > 3 billion cells, see **Fig. 4**), applying a binary decision rule with a threshold of 0.05 percentage points per hour for dSGI revealed that only the vessel harvested on Day 7 would have satisfied the criteria for harvest *a priori* (dSGI dropping below the dotted line threshold in Fig 5b) – demonstrating the potential utility of dSGI as a real-time harvest indicator. Therefore, users of the Skroot sensor system may perform application-specific cell culture runs to determine a suitable dSGI decision threshold for harvest readiness.

In our experiment, both the continuous and benchtop modes would have indicated that the manufacturing process was harvest ready approximately 24 hours prior to Day 7. The continuous dSGI crossed the dSGI threshold earlier as the benchtop dSGI was constrained by the discrete sampling intervals. Nonetheless, the ability of the benchtop reader to
support harvest decisions with limited data is noteworthy enabling a simple yet effective “go/no-go” harvest indicator (Supplement 7).

However, the higher temporal resolution of continuous mode could enable users to predict the onset of the harvest window further advance, thereby potentially enhancing the operational efficiency of the manufacturing process (Supplement 8).

### Impact of SMART Sensor on Production Costs

The sensors described herein directly affect the variable costs for each dose produced by reducing the amount of direct labor time of intermittent sampling and end cell count present in many use cases. The sensors can also detect expansions that are not growing early for time-effective restarts. Scaling to >100,000 doses must be achieved to reduce the burden of overhead costs currently dominating the cost structure of these therapies at clinical and commercial scale. The logic is straightforward: the facilities and equipment are expensive, and a large workforce is required, but only a limited quantity of products can be produced in the current paradigm; with high overhead costs and low throughput, the costs required to run a high functioning cell therapy manufacturing operation distributed on a per drug product production basis at both clinical and commercial scale are leading to several company failures despite promising clinical data and some commercially approved therapies are struggling to maintain a viable business model (see Supplement 9 for further discussion). Until we can reach high production scale, overhead costs will continue to dominate the cost structure of producing these therapies and therefore cost of materials, consumables, and reagents are negligible and insignificant when it comes to lowering the costs of these drug products to the patient. Therefore, any improvements to the manufacturing process that reduces the amount of direct labor time per drug product and can increase throughput on a given production line, will contribute to significant reduction in per dose costs. A G-Rex based manufacturing process with a highly functional manufacturing operation can achieve the scales contemplated herein and the SMART sensor has the potential to further augment G-Rex simplicity and scalability through a reduction in direct labor, while enhancing the manufacturer’s ability to monitor and optimize the manufacturing process.

## Conclusions

Herein, we have shown the application of the SMART sensor platform to the G-Rex production system for highly scalable, cost-effective production of cell and gene modified cell therapies. In this research, the sensor was optimized for scaled production and used in simulated clinical applications. It was validated with cytotoxicity testing, aging studies, and through demonstration with donor PBMCs. The cell work showed the ability to detect different growth rates, provide an indication for harvest timing, and the potential to eliminate the need of a cell count prior to cryostorage of drug product. We also demonstrated the reliability of this method as a surrogate measure for cell counting and periodic metabolite measurements thereby establishing its utility in eliminating periodic sampling for those applications that may require it while providing an option for continuous monitoring of cell expansion for G-Rex applications where this feature may be of use.

Beyond the adoption of these new sensor systems, this work also presents a compelling call to continue standardizing and improving highly scalable parallel production methods, the central focus of G-Rex based cell therapy manufacturing.

## Supporting information

Supplement Sections

## Funding

This work was supported in part by National Science Foundation SBIR Grant Award# 2025552

## Competing Interests

The SMART G-Rex and accompanying reader are currently being developed as a commercial product to be sold through Wilson Wolf Manufacturing LLC. The author team hold patents in G-Rex and SMART sensors and readers.

## Acknowledgements

We would like to thank Eric Smith, Andrew Lewis, and Nathan Dunkerley for early discussions on the SMART G-Rex concept.

## Notes

### Competing Interest Statement

The authors from Skroot are authors on patents pertaining to the use of SMART sensors for cell therapy manufacture. The authors from Wilson Wolf are authors on patents pertaining to the use of G-Rex for the manufacturing of cell therapies. The joint team seeks to commercialize this SMART G-Rex platform technology and have financial interest in its success. The authors were objective in their scientific approach and present all data and analysis such that others can reproduce this work.

