## Supplement Sections for "SMART Sensors for Cell Growth Monitoring in Closed System G-Rex for Scalable, Cost-Conscious Cell and Gene Therapy Manufacturing"

### Supplement 1: SMART Sensor Response to T Cell-Secreted Signaling Molecules

T cells are known to secrete lipid mediators derived from polyunsaturated fatty acids (PUFAs), specifically eicosanoids and their derivatives, such as prostanoids (like $PGE_2$) and leukotrienes. These signaling lipids function as key modulators of immune responses, influencing T cell functions like migration and differentiation. Arachidonic acid (AA), an omega-6 PUFA that is essential for cell membrane structure and a source of energy, serves as a precursor to eicosanoids.

To validate the SMART sensor transduction method for T cells, an experiment was performed using two 35 mm petri dishes (control and experimental) with embedded SMART sensors. Each dish was initially filled with 3 mL phosphate-buffered saline (PBS) to establish a 5 hour long baseline response using SMART readers in continuous mode. Subsequently, 0.25 mL of an AA solution (100 mg AA dissolved in 1 mL ethanol) was added to the experimental petri dish, while 0.25 mL of pure ethanol was added to the control. As shown in **Figs. S1-1a** and **S1-1b**, AA addition produced a distinct change in sensor response compared to the control. A second addition of 0.75 mL of the AA solution to the experimental dish, paired with 0.75 mL ethanol to the control, induced a further pronounced response, demonstrating the sensitivity of the SMART sensor transduction method to AA, a precursor of T cell-derived lipid mediators.

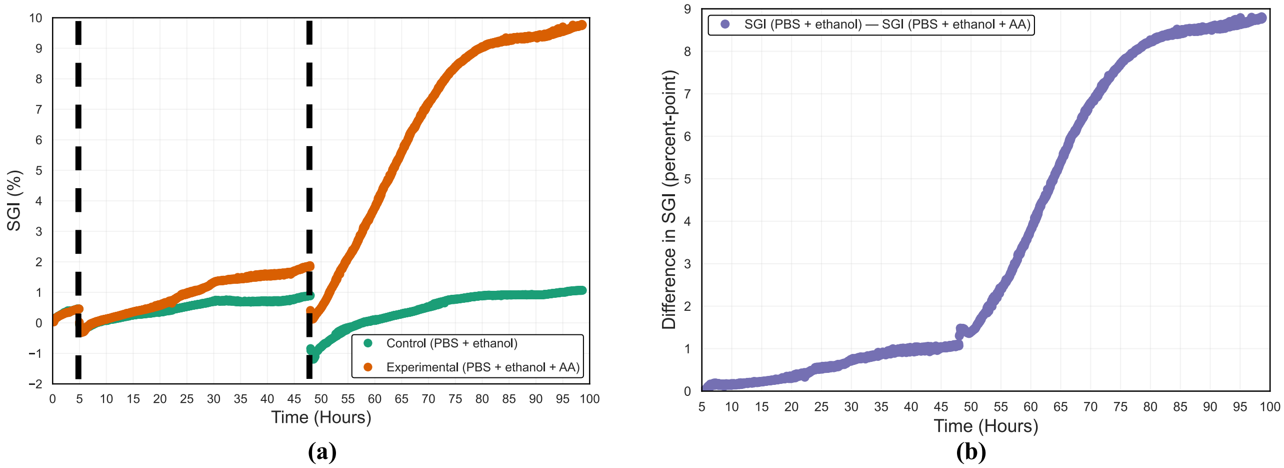

**Figure S1-1** – (a) SMART sensor response for control and experimental petri dishes showing clear effect of Arachidonic Acid (AA). The two vertical dashed black lines correspond to instances of addition of chemicals to the dishes. (b) Difference in sensor response between two dishes, from the point of first chemical addition, further emphasizing the transduction of the sensor by AA. Note that this plotted difference is simply one SGI plot subtracted from another, and is different from the quantity dSGI, which indicates the finite time derivative of SGI.

### Supplement 2: Effect of Kerf on Sensor Response

In this work, the PET layer in the sensor stack was laser cut to form high-precision air voids that enabled the transduction membrane (softened in response to secondary metabolites in the cell culture) to collapse and fill the voids, thereby altering the resonant sensor's response, quantified in terms of the Skroot Growth Index (SGI). The cross section stack of the sensor is revisited in **Fig. S2-1a**.

It was observed that changing the kerf setting on the laser cutter, which controls the width of the PET material removed, significantly affected the saturation behavior of the SMART sensor. In particular, a wider kerf resulted in a greater overall shift in SGI and a faster saturation to the final value. **Figs. S2-1b** and **S2-1c** illustrate this effect for *E. coli* and CAR-T cell cultures, respectively.

Thus, kerf width represents a useful design parameter for tuning sensor response to track cell growth metrics of interest. Herein, a nominal kerf width of 300 μm was ultimately selected as a trade-off between the sensor's ability to track lactate (see the 350 μm trace in **Fig S2-1c**) and the tolerance limitations of the third-party sensor manufacturer.

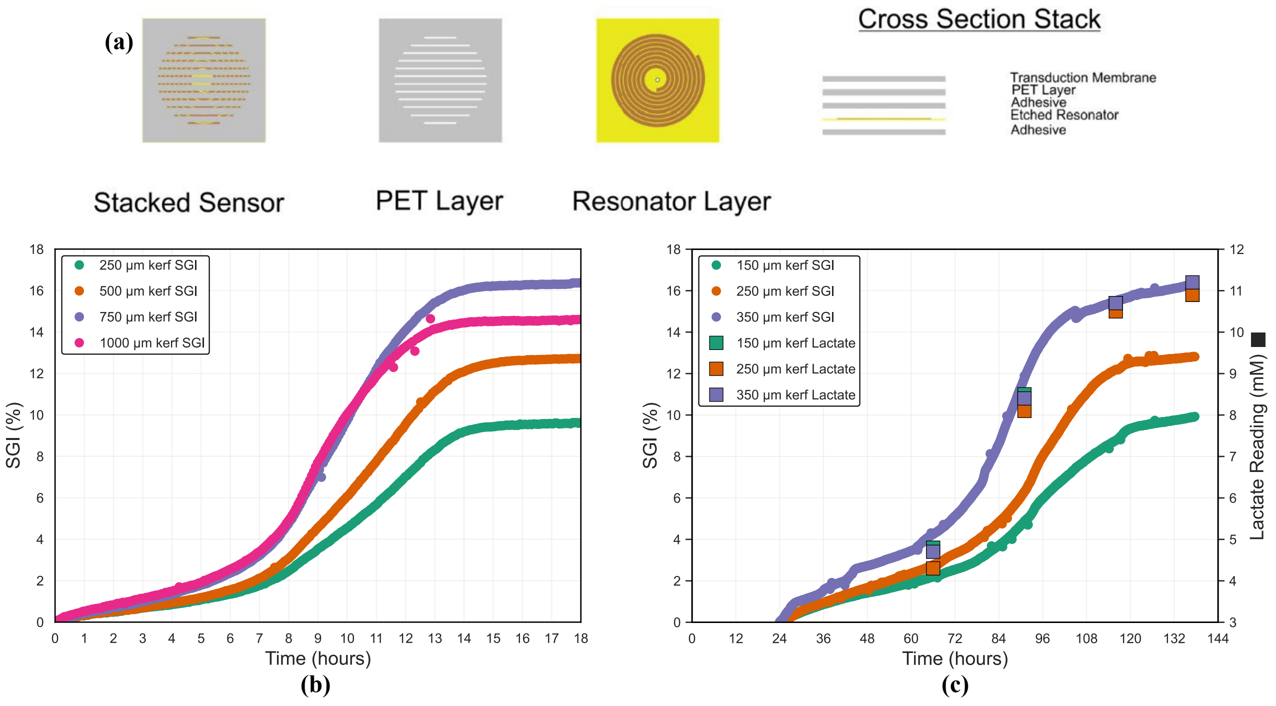

**Figure S2-1** – (a) Cross-sectional stack of the SMART sensor, highlighting the cut channels in the PET layer. The channel width (kerf) is a tunable design parameter that modulates sensor response. This effect is demonstrated in (b) E. coli and (c) CAR-T cultures, where larger kerf widths result in greater SGI shifts and faster saturation dynamics.

### Supplement 3: Cytotoxicity Testing and Accelerated Aging Testing for Third-party Sensors

*Cytotoxicity Testing*

A cytotoxicity assay was conducted on the SMART sensors adhered to polycarbonate caps (same material as the G-Rex 100M body) by NAMSA. The study was performed in accordance with ISO 10993-5 and BS EN ISO 10993-5, which specify tests for in vitro cytotoxicity of medical devices, and adhered to FDA Good Laboratory Practice (GLP) Regulations (21 CFR, Part 58)

For the evaluation, a single preparation of the Cap with Sensor was extracted in Eagle's Minimal Essential Medium supplemented with 5% Fetal Bovine Serum (EMEM 5% FBS) at 37 °C for 24 hours with continuous agitation. The chosen extraction vehicle was designed to optimize the extraction of both polar and non-polar components from the test article. Triplicate monolayers of L-929 mouse fibroblast cells, a historically used and established cell line sensitive to cytotoxic leachates, were then dosed with the prepared extract. These cell monolayers were subsequently incubated at 37 °C in 5% $CO_2$ for 72 hours, after which they were microscopically examined for any signs of abnormal cell morphology or cellular degeneration. Reactivity was graded on a scale from 0 (none) to 4 (severe).

The validity of the assay was confirmed by the performance of the control articles. The negative control (High density polyethylene) and the vehicle control (EMEM 5% FBS) both exhibited no reactivity, receiving a grade of 0. Conversely, the positive control (0.1% Zinc diethyldithiocarbamate) demonstrated severe cytotoxicity, consistently scoring a grade of 4.

The test article extract showed no evidence of causing cell lysis or toxicity in any of the test wells at 72 hours. All three replicates of the received a reactivity grade of 0 (None). This result indicates that the test article extract met the requirements of the test, as its biological response was well within the acceptable limit of less than or equal to a grade 2 (mild reactivity). The conclusions drawn are specific to the test article evaluated in this study.

*Accelerated Aging Testing*

Four SMART sensors were applied to each G-Rex 100M device which were then gamma sterilized. Subsequently, the G-Rex 100M devices were split into four different aging groups – Year 0 through Year 3. While the Year 0 devices were stored at room temperature, the other groups thermally aged inside a Fisherbrand™ Gravity Oven at Skroot Laboratory, Inc. for a predetermined time for each aging group calculated using Accelerated Aging Theory[1]:

$$t_{accelerated} = \frac{t_{desired}}{Q_{10}^{(T_{elevated}-T_{ambient})/10}} \quad \text{(S3-1)}$$

where $t_{accelerated}$ and $t_{desired}$ respectively denote, the accelerated aging and desired real time aging time durations, $Q_{10}$ is a constant with value 2.0, while $T_{elevated}$ and $T_{ambient}$ are the elevated and ambient temperatures respectively. Using $T_{elevated}$ as 59 °C and $T_{ambient}$ as 22 °C, resulted in the following accelerated aging time durations for each aging group.

**Table S3-1** – Sensor Aging Group and Accelerated Aging Time Duration for Each G-Rex 100M Used for Aging Test

| G-Rex 100M Identifier | No. of Sensors | Sensor Aging Group (Years) | Accelerated Aging Time Duration (Days) |
|---|---|---|---|
| V-0144 | 4 | 0 | 0 |
| V-0147 | 4 | 1 | 28 |
| V-0150 | 4 | 2 | 56 |
| V-0153 | 4 | 3 | 84 |

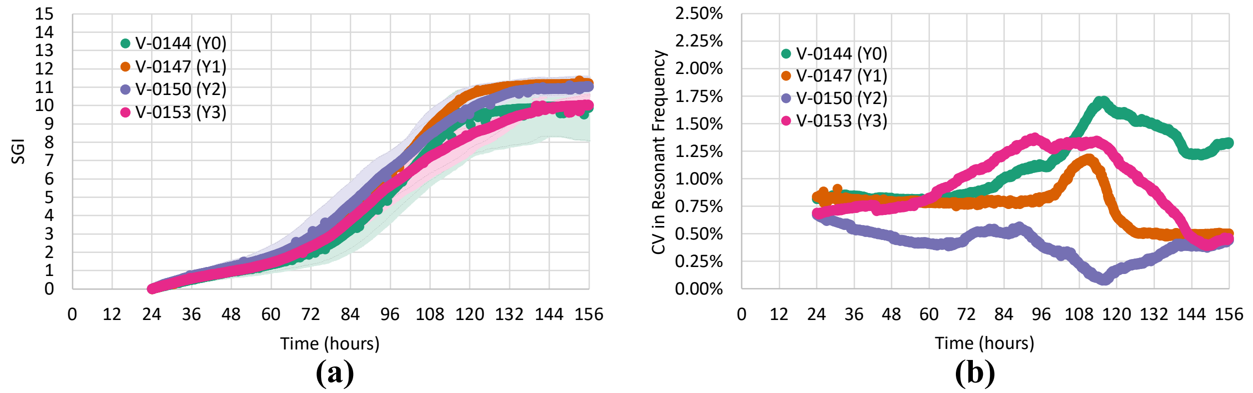

**Figure S3-1** – (a) Mean SGI with min and max error bars (denoted by tinting the primary color) for G-Rex 100M devices aged 0 through 4 years, (b) coefficient of variation (CV) in resonant frequency measurements for the 4 G-Rex devices.

[1] Hemmerich, K. J., "General Aging Theory and Simplified Protocol for Accelerated Aging of Medical Devices," Medical Plastics and Biomaterials, July/August 1998, pp. 16–23.

Upon completion of accelerated aging for each aging group, the G-Rex 100M devices were stored at room temperature until all groups were done aging.

The cell culture test involved placement of the G-Rex 100Ms on Skroot readers. Cell seeding was done on Day 0 using the medium growth condition (as defined in the main text) from the same PBMC bag for all G-Rex 100M devices.

Aging over a 3-year period did not adversely impact sensor reliability, cell growth tracking, or data quality (**Fig. S3-1 a, b**). These findings support the use of Skroot sensors for long-term storage without compromising downstream biomanufacturing performance.

### Supplement 4: Analysis of Raw Reader Signal to Compute SGI

For every frequency sweep, the reader hardware interrogates the sensor-under-test wirelessly at each frequency point in the sweep and outputs a digitally converted DC voltage value. This DC voltage versus interrogation frequency curve peaks when the interrogation frequency is equal to the resonant frequency of the sensor. To account for the reader's intrinsic frequency response and cancel off any systematic errors, each frequency sweep of the sensor is calibrated against a reader-only calibration response (no sensor present) done before the start of each experiment. **Fig. S-4-1a** demonstrates a sample calibration response, while **Fig. S-4-1b** shows the calibrated sensor response (signal strength) for the first and last frequency sweeps from a 7-day long cell culture run with a SMART sensor. The calibrated sensor response at each frequency point ($f$) is calculated in software as the following and expressed in percentage form:

$$V_{sensor,calibrated}(f) = \left(\frac{V_{sensor,raw}(f)}{V_{calibration}(f)} - 1\right) \tag{S3-1}$$

where $V_{sensor,calibrated}$ and $V_{sensor,raw}$ represent the calibrated and raw sensor outputs, respectively, with $V_{calibration}$ being the reader-only calibration response. The calibrated sensor output is a measure of the percent change in the strength of the output signal influenced by the sensor on top of the baseline calibration response, and is thus termed as signal strength. Major factors which impact signal strength include the coupling coefficient between the reader antenna and the conductive losses in the media and a lower signal strength makes it harder to detect the resonant peak due to the sensor.

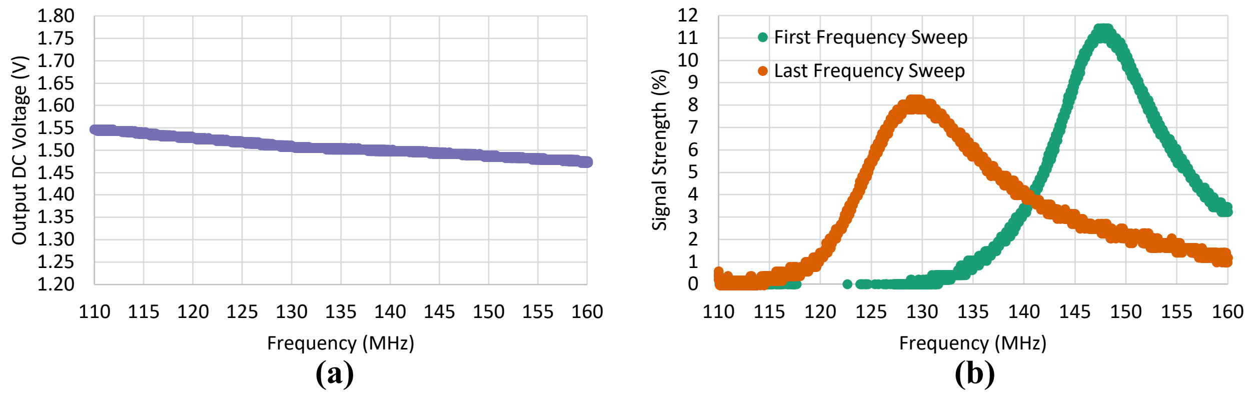

**Figure S4-1** – (a) Calibration response of reader hardware showing output DC voltage over an entire frequency sweep, (b) calibrated sensor response from the first and last frequency sweep of a 7-day long cell culture run clearly showing the shift in resonant (peak) frequency.

The peak itself is extracted by fitting a 500-point Gaussian curve around the absolute maximum and the corresponding interrogation frequency point is regarded as the sensor resonant frequency, from which the software can readily calculate SGI using equation (1) from the main text. Using this formulation, in **Fig. S-4-1b**, given that the first and last frequency sweeps yield resonant frequencies of 147.29 MHz and 128.86 MHz respectively, the total SGI shift (assuming an equilibration time of 0) is $(147.29 - 128.86)/128.86 = 12.51\%$.

Finally, a Savitzky-Golay digital filter[2] is to smooth the real-time SGI curve.

[2] Schafer, R. W., "What Is a Savitzky-Golay Filter? [Lecture Notes]," *IEEE Signal Processing Magazine*, vol. 28, no. 4, 2011, pp. 111–117.

### Supplement 5: Effect of Reader Temperature on Cell Growth

During the early development phase of the Skroot platform, a commercial portable vector network analyzer (VNA) was used to interrogate the SMART sensor. However, it was observed that G-Rex devices with active readers resulted in a lower harvest cell count compared to control vessels not using Skroot technology (**Fig. S5-1a**), without an observable change in cell viability (**Fig. S5-1b**). Temperature measurements (**Fig. S5-1c**) showed a ~1 °C increase when using Skroot readers leading to the hypothesis that the heat generated by the VNA was increasing the temperature and hindering cell growth.

This hypothesis led us to swap the VNA with a custom-designed reader which had careful hardware and firmware design to manage the heat generated by the system. In particular,

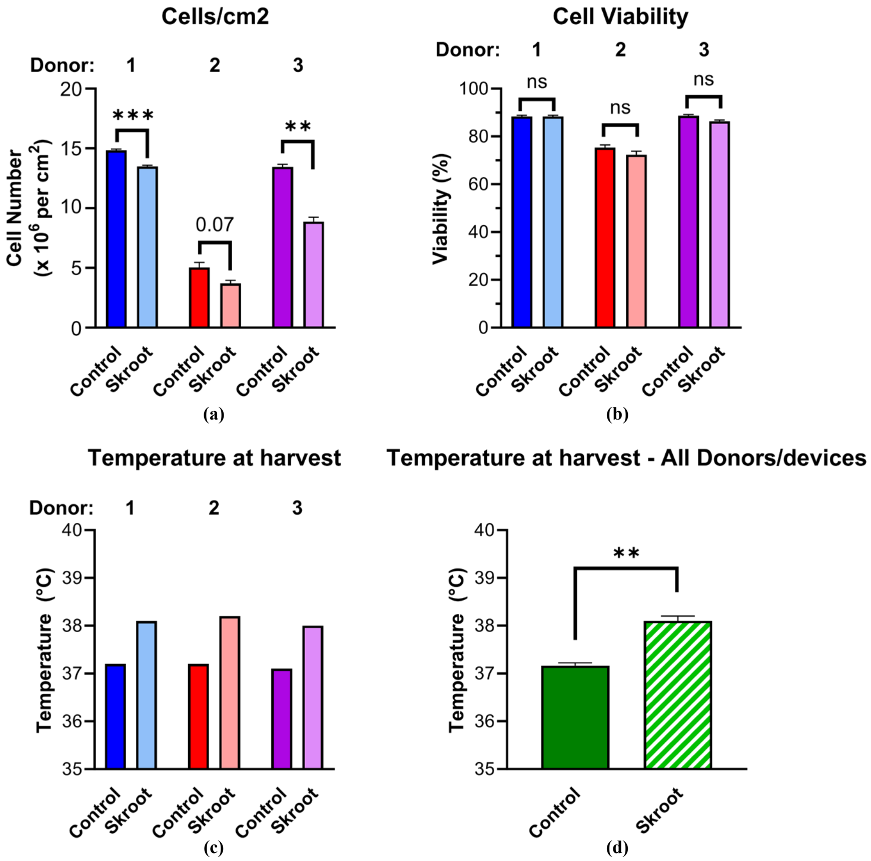

**Figure S5-1** – (a) Comparison of cell count in control versus early-stage Skroot G-Rex devices (with active VNA readers) showing lower harvest cell count across three cell cultures with independent donors with the early-stage Skroot platform (b) without a significant drop in viability. (c) The temperature of the vessel upon harvest was consistently greater by ~1 °C for devices using the Skroot platform, (d) characterized further by the average temperature measurement.

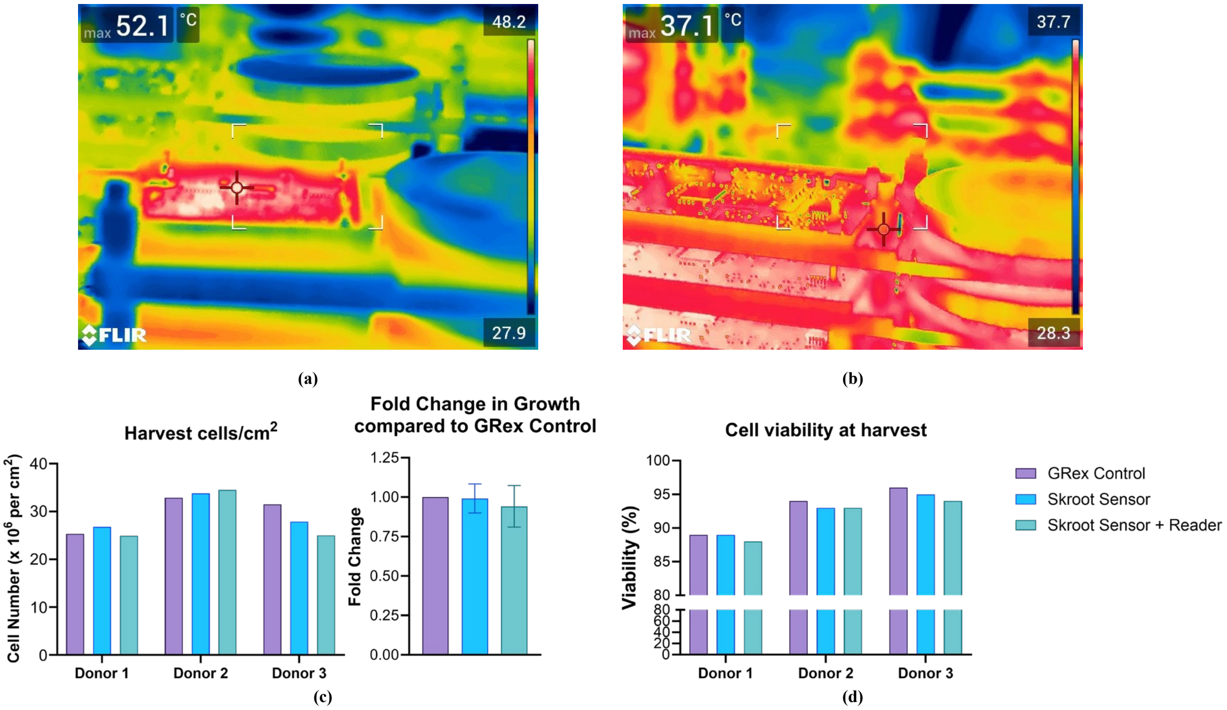

**Figure S5-2** – Thermal images to compare the temperature of the (a) commercial VNA reader and (b) custom-designed reader showing that the custom-designed reader did not heat up like the VNA over the course of its operation. Cell testing with the new custom-designed reader demonstrated no statistically significant difference from control G-Rex devices in terms of (c) cell count and (d) cell viability upon harvest.

special attention was paid to the power consumption of electronic components selected for the design, while the firmware on the custom-designed reader ensured that the hardware was put to sleep when not actively scanning sensors which stopped the incubator interior from heating up due to reader operation. Thermal images comparing the VNA (**Fig. S5-2a**) and the custom-designed reader (**Fig. S5-2b**) showed that the hottest part of the latter was significantly lower (and approximately the ambient incubator temperature). Finally, cell testing with the new readers showed no statistical difference (using two-way ANOVA with Bonferroni's multiple comparisons testing) in harvest cell count (**Fig. S5-2c**) and viability compared to control G-Rex devices (**Fig. S5-2d**).

### Supplement 6: SMART Platform Validation Across Cell Types

The SMART sensor platform was validated across multiple cell types, including fast-growing bacteria and yeast as well as slower-growing mammalian cells such as CHO and K-562 in addition to CAR-T as discussed in the main text. The sensors successfully tracked cell growth in both static and dynamic culture vessels, with sensor outputs benchmarked against standard manual sampling parameters (**Figs. S6-1a–c**). Furthermore, the sensors demonstrated the ability to noninvasively detect aberrant cell culture conditions such as contamination. As shown in **Fig. S6-1d**, contamination induced a rapid increase in SGI followed by early signal saturation, highlighting the platform's potential for real-time contamination detection. Future work analyzing the derivative of SGI may enable development of a robust, continuous contamination monitoring tool.

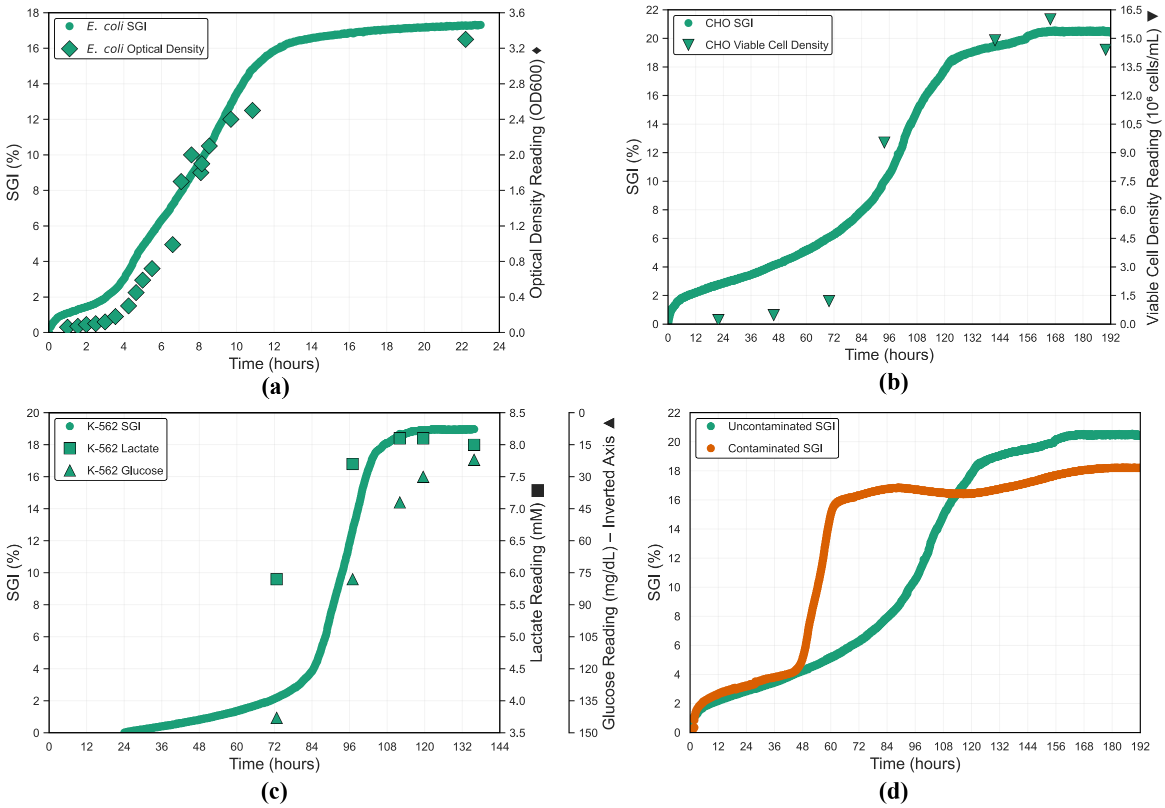

**Figure S6-1** – SMART sensor response for three representative cell types: (a) E. coli cultured in a 250 mL shake flask, with SGI compared to manually sampled optical density (OD600); (b) CHO cells cultured in a 250 mL shake flask, with SGI compared to manually sampled viable cell density; (c) K-562 cells cultured in a G-Rex 100M, with SGI compared to manually sampled lactate and glucose concentrations (glucose plotted with an inverted right axis); along with (d) comparison of SMART sensor response in uncontaminated and contaminated CHO cell cultures grown in 250 mL shake flasks. Contamination resulted in a rapid rise in SGI followed by early saturation.

### Supplement 7: Harvest Indicator Validation

Herein we provide an example of a simple binary “go/no-go” (ready for harvest/not ready for harvest) indicator based on benchtop reader data. An experiment was conducted at CellReady, LLC involving 10 G-Rex 100M devices – all seeded on the same day (Day 0),

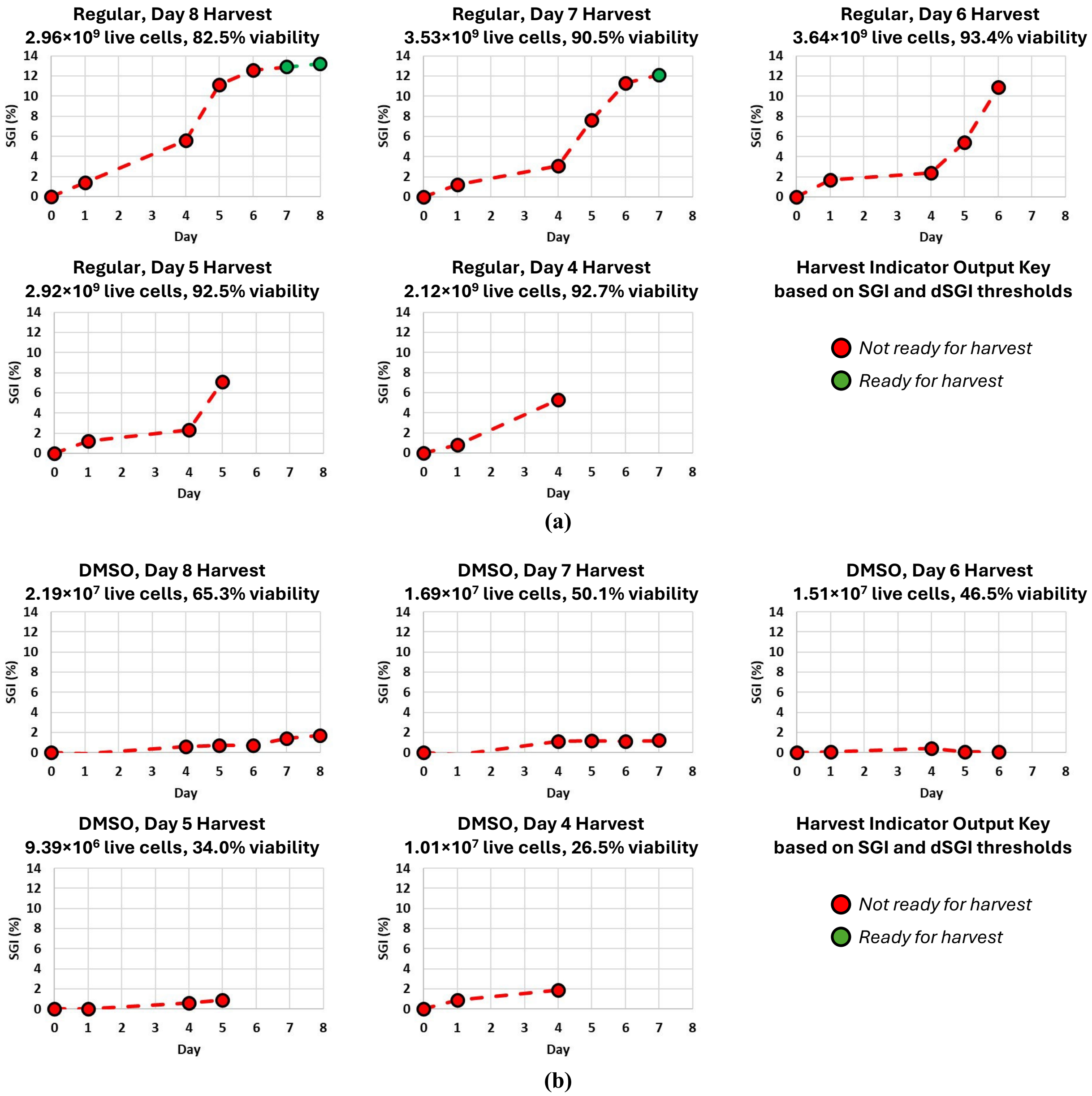

**Figure S8-1** – (a) SMART sensor response using benchtop reader for 5 G-Rex 100M devices that were identically seeded with 50 million cells on the same day (Day 0), activated together, and harvested on Days 8, 7, 6, 5, and 4 along with the binary “go/no-go” output of the harvest indicator. (b) The same set of plots for 5 G-Rex 100M devices that were additionally spiked with 10 mL 2.5% DMSO solution to stunt cell growth. Corresponding to each vessel, also shown are terminal live cell count and viability. It is evident that the harvest indicator successfully recommended harvest in the window with the highest live cell count.

identically, with 50 million cells from a PBMC bag. The 10 vessels were equally split into two groups – one which was allowed to grow using CellReady's standard protocol while the other had each vessel spiked with 10 mL of a 2.5% dimethyl sulfoxide (DMSO) solution to stunt cell growth. For each group, the 5 vessels were harvested successively on Days 4 through 8. On each day, SMART reader (in benchtop mode) scans were taken along with manual samples for lactate and glucose for all active vessels, while terminal cell count and viability were recorded upon harvest. The reader implemented a real-time harvest indicator algorithm based on a decision rule using thresholds of SGI > 7% and dSGI < 0.05 percentage points per hour (as described in **Fig. 5**). The performance of the harvest indicator was validated against the terminal live cell count and it was found that the indicator was successfully triggered only when live cell count was the highest (**Fig. S8-1a**). The only scope of improvement was early detection of the regular vessel harvested on Day 6 – finer resolution provided by more frequent/continuous reader scan could have facilitated it. Nevertheless, it can be inferred from the Day 7 harvest vessel, that had the Day 6 harvest vessel been run for an extra day, the harvest indicator would have been triggered with the live cell count still sufficiently high.

Moreover, with the stunted growth condition (DMSO), no false positives were observed – further corroborating the effectiveness of the harvest indicator. Additionally, the DMSO vessels also point toward the possibility of implementing a threshold for early detection of slow or abnormal growth which could enable a manufacturing facility to intervene and course correct.

### Supplement 8: Harvest Predictor Proof-of-Concept

This supplement shows the proof-of-concept for developing a real-time harvest predictor based on continuous SMART sensor data. **Fig. S8-1** illustrates SGI and dSGI for a K562 culture demonstrated above. The data received up to time = 100 h was used to predict when SGI would saturate in terms of dSGI falling under a predetermined threshold (similar to the harvest indicator discussed in the main text). For this particular use case, a Gaussian fit around the critical growth rate (peak in dSGI) was used with a dSGI threshold of 0.05 percent-point/hour, resulting in a saturation prediction 8 hours in advance. The sensor response post 100 hours shows the predicted harvest window indeed corresponded with the beginning of SGI saturation. The prediction would continue to be refined as more incoming data was processed in real-time beyond the 100 hour mark.

Future research would involve incorporating a robust machine learning algorithm, in accordance with application-specific calibration inputs, to accurately predict harvest readiness in mammalian cells at least 24 hours ahead of time.

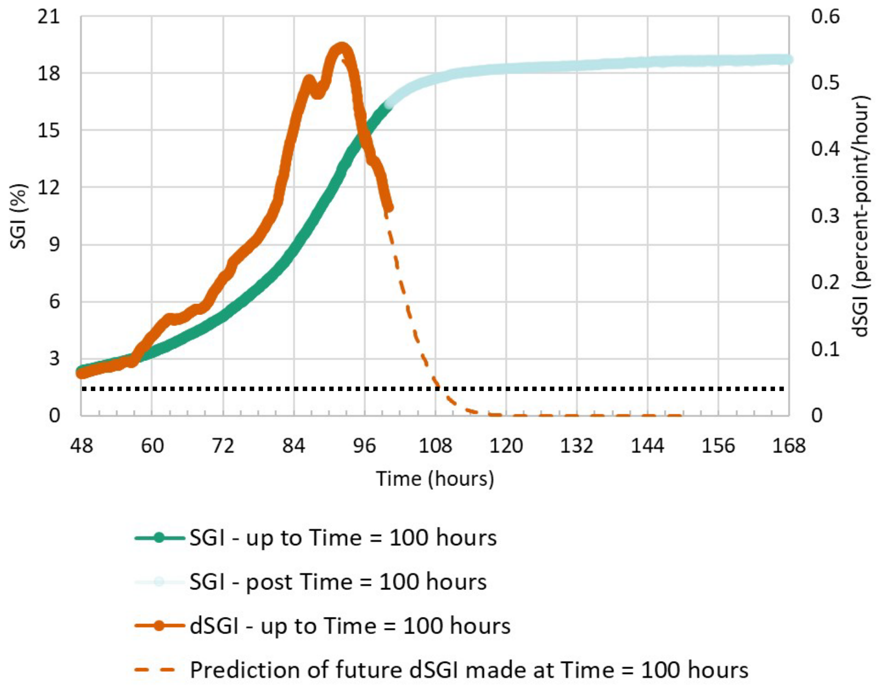

**Figure S8-1** – SGI and dSGI for K562 culture in T flask. Data up to 100 hours was used to predict dSGI falling lower than a predetermined threshold of 0.5 percent-points/hour. The SGI data post 100 hours demonstrate the accuracy of the prediction.

### Supplement 9: Cost Discussion

Systems such as the Miltenyi Biotec Prodigy, Lonza Cocoon, Ori Biotech IRO, or Cellares Cell Shuttle, have promised a reduction in cell therapy production costs through "automated" process steps, but these technologies hit a fundamental limit due to their inherent cycle time limitations as well as requiring complex and expensive consumables and associated software systems without commensurate reductions in human resource requirements. Each product is made one at a time and the significant capital equipment and valuable production floor space is tied up until the therapy is fully complete (usually around one to two weeks per product).

A cost study[3] of CAR-T produced at an academic hospital using a proven system (Prodigy) demonstrates this limit by estimating maximum annual production of only 54 doses using three Prodigy systems in a 215 $ft^2$ cGMP space. The study estimates a cost of goods (COGs) of $60,000 per dose at full production, but this likely neglects the full burden of overhead, and the costs of maintaining GMP compliant inventory management systems, a compliant and nimble Quality Management System, maintaining trained manufacturing and QC personnel and fully ignores the high set-up and operating costs of purchasing the dedicated equipment, building out dedicated cGMP space, dedicated QC labs, and maintaining both fully staffed with dedicated and trained personnel.

Moreover, these academic price assessments likely underestimate overhead costs because manufacturing personnel are often post-docs or researchers and their costs to the academic hospital can be subsidized by existing grants unrelated to the production of the clinical drug products in their cGMP space. Additionally, equipment, space, and personnel can be distributed over a greater number of clinical manufacturing projects because most of these clinical products produced in an academic hospital setting have smaller phase 1 patient populations. Therefore, the GMP space can be used for multiple clinical product manufacturing, however, this quickly becomes untenable if clinical production demand for any of the products increases. When this happens, the problem becomes an inability to meet production demand. Interestingly, this is a small-scale version of the problem facing the entire industry.

The industry is struggling to manufacture enough product to treat the addressable market. An analysis of the problem on a small scale in an academic institute will highlight some important considerations for the entire field. The estimated 54 doses per year using 3 Prodigy systems likely overestimates true capacity since the manufacturing line cannot be balanced, because the cycle time on the instrument is inherently tied to the entire

---

[3] https://pubmed.ncbi.nlm.nih.gov/32535920/

production process, and demand is never perfectly balanced. When production demand for a particular clinical candidate reaches this threshold, in our view, the sponsor has a few options: 1) Ask the academic institute to increase capacity in its existing location, 2) add production capacity by setting up manufacturing at another academic hospital with the same or similar GMP operation, 3) move to a centralized CDMO with greater production capacity, or 4) take operations in house to a centralized dedicated facility funded by the sponsor.

All these options come with cost tradeoffs and operational risks to the sponsor. While recognizing that the following analysis contains forward-looking statements and assumptions that should be challenged, we attempt to critically evaluate the options available to the sponsor:

> Option 1 will be a difficult proposition because the hospital or academic center will require additional demand from other sponsors in order to justify the additional investment. If the demand exists, it may ultimately lead to a similar bottleneck in production capacity; on the other hand, there will be risk to the hospital or academic center that sponsor funding dries up if positive clinical data are not obtained, which would risk underutilization of resources and less ability to distribute overheads from grant funding. The institute would be forced to continuously court new customers. In today's funding environment, it is likely difficult to achieve alignment with the academic center on whether the investment and additional overhead cost is worth the risk.
>
> Option 2 will require significant coordination between the sponsoring entity and the new entity and the bottleneck in capacity may quickly be reached again. This would be a version of the often-cited point of care manufacturing or modular manufacturing model. Some governments have provided guidance on this model, for instance the Medicines and Healthcare products Regulatory Agency of the UK and Northern Ireland recently published guidance on Point of Care and Modular Manufacturing of ATMPs and other products[4]. The Guidelines make it clear that license holders must ensure the various sites manufacturing under the license comply with current good manufacturing practice and must maintain staff, premises, and equipment necessary for manufacture of the products in accordance with the master file submission. Additionally, the license holders must maintain supervision and control of manufacturing changes, corrective and preventive actions, non-conformance reporting, and release activities. In other words, to

[4] https://www.gov.uk/government/publications/human-medicines-modular-manufacture-and-point-of-care-regulations-2025-overview/human-medicines-modular-manufacture-and-point-of-care-regulations-2025-overview

employ this model, the adopting sites must have a Quality Management System and ideally all sites would have the same Quality Management System otherwise maintaining adequate control will become overwhelming and impractical. Operating each manufacturing site under a different Quality Management System would be comparable to a large franchise allowing each of its franchisees to use a different operating system and kitchen layout. Scaling this model to large patient populations will require a significant number of sites and if this is done in collaboration with hospitals rather than as sponsor owned entities operating completely independently, then there are likely to be resource constraints, contract disputes, and strategic misalignment as hospitals will seek to offer care to patients outside of the scope of a manufacturing license holder's indication and may prioritize other projects at the expense of the sponsor's interests. The issue is further exacerbated when additional clinical indications are added, and new treatments become available. Therefore, for a more distributed model of ATMP manufacture, it may be more practical to deploy modular manufacturing where the sponsor entity fully owns and controls the site, however the complexity of managing multiple sites rather than a single site remains, and there are additional capital expenditures and opportunity costs qualifying the new facility and running adequate comparability studies and technology transfers from the main site to a new modular manufacturing site.

Option 3 will enable more production capacity, but there is still a large upfront technology transfer fee and the likely increase in costs caused by higher cleanroom reservation fees at CDMOs required to enable scheduling of patient production and secure manufacturing capacity at the site. CDMOs are also subject to high levels of production personnel turnover and funding issues. Lastly, since the industry does not have sufficiently standardized production operations or manufacturing platforms, the current CDMO model requires that the CDMO maintain various equipment platforms so that they can have access to a wider customer base. The upside to the CDMO being that any sponsor independent of their manufacturing platform is a potential customer, but the downside is that they need to maintain more suite space than they otherwise would have to while maintaining multiple manufacturing platforms, most of which take up significant space. Inherently, when the CDMO onboards multiple customers with different platforms of choice, the CDMO needs to charge high reservation fees to facilitate scheduling of patient slots with the sponsor because the instruments tie up the space and scheduling is often unpredictable. Thus, the CDMO is forced to allow equipment and space to remain idle on behalf of their clients in order to accommodate unpredictable patient manufacturing demand.

Notably, most, if not all these problems are eliminated if a G-Rex based manufacturing platform is adopted by the industry and by CDMOs. The G-Rex based CDMO will have access to a large customer base, their operators will all be familiar with closed system G-Rex operations, and they'll be able to distribute overhead costs across a larger number of sponsors and patients. Reservation fees may be foregone entirely provided the CDMO maintains the equipment necessary to process starting material ahead of the projected demand. Eliminating or reducing reservation fees will make for an attractive pricing strategy to potential customers and therefore more likely to win customer business. Until more CDMOs recognize the significant business advantages that a G-Rex based CGT production operation presents them, most sponsors will face the above-mentioned financial and operational issues by choosing this Option 3.

Option 4 provides the most control to the Sponsor theoretically, but it comes with the highest upfront capital expenditures and will also come with higher cash burn rates because the sponsor will be forced to pay larger overheads to staff a manufacturing facility. Using Prodigy as an example, based on the projection of 54 patients per three Prodigy, a Sponsor would need to purchase 6 Prodigy just to service an annual patient population of around 100. To scale this to meaningful patient populations in a centralized facility would require 10x this initial capital expenditure and even then, it would be difficult to balance the manufacturing line with the unpredictable production demand.

The overarching goal of the sponsoring entities of these early-stage clinical assets should be to obtain as much clinical data as possible without requiring significant cash outlays so that less investment money is required to achieve clinically meaningful data. This would reduce the risks to investors associated with building out an entire manufacturing operation and staffing it with appropriately qualified personnel should that be needed later. In general, the scale of clinical demand for phase 2 and phase 3 clinical trials (i.e., 50-300 patients) would not normally justify the upfront capital expenditures. Unfortunately, this is an option that many companies have taken, and it has led to unsustainable operating costs that are not economically feasible for the sponsor. It does not make sense to operate a facility that costs $10M-20M per year annually and often much more before finding out if the drug product candidate is likely to succeed commercially by generating the clinical data to de-risk further investment.   Once again, notably, these issues are mitigated with a G-Rex based production platform because the upfront capital equipment investment is minimized and the space requirements are significantly diminished.

What is common to all of these options is a critical need for standardized CGT manufacturing operations based around a technology that does not unnecessarily tie up capital equipment for extended periods of time. The SMART G-Rex further advances this goal of standardized protocols with less intervention points thereby increasing throughput and decreasing the direct labor cost per manufactured dose.

By focusing solely on semi-automated and non-standardized production systems that are successive in nature (or modestly parallel in the case of the Cell Shuttle), that tie up valuable space, equipment, and personnel time, pharmaceutical companies, and academic institutes alike, will continue to struggle to make CAR-T manufacturing a long-term viable option. More importantly, they will continue to struggle to deliver the quantity of drug product needed to meet the total patient population and deliver the true promise of these therapies on a large scale.
